# A scalable filtration-based method for isolating exomeres and other nanoscale extracellular particles

**DOI:** 10.1101/2025.10.23.684032

**Authors:** Thomas Scarborough, Yuki Kawai-Harada, Olivia Brennan, Christina Chan, Masako Harada, S. Patrick Walton

## Abstract

Extracellular particles, including extracellular vesicles (EVs) and non-vesicular extracellular particles (NVEPs), enable intercellular communication by transferring regulatory miRNAs and other biomolecules. While EVs have been studied for drug delivery, NVEPs remain relatively unexplored. Exomeres, a recently discovered class of NVEPs enriched in RNAi proteins, preferentially carry miRNAs and deliver them to cells more effectively than EVs, underscoring their potential as vehicles for therapeutic RNAs. One current limitation to studying and applying exomeres for therapeutic RNA delivery is the shortage of scalable, cost-effective, and rapid isolation methods. Here, we investigated whether tangential flow filtration (TFF), a common bioseparation approach that separates species by size, would effectively isolate exomeres from conditioned media with comparable purity and identity to those isolated by differential ultracentrifugation. TFF successfully isolated exomeres that were enriched in RNAi components including AGO2, HSP90AB1, and a unique set of miRNAs not abundant in EVs. Remarkably, exomere miRNAs were resistant to nuclease degradation even after treatment with protease and surfactant, suggesting that exomeres are highly stable, non-vesicular complexes with potentially extended circulating half-lives. Together, our results establish TFF as an efficient method for isolating exomeres, and further demonstrate that TFF could be applied in a bioprocess for exomere-based RNA therapeutics production. This study is also the first to demonstrate that exomere miRNAs are highly resistant to nuclease degradation, suggesting that exomeres could complement and potentially outperform current clinical standards for RNA delivery.

## Introduction

It is now well-established that intercellular communication is mediated by a variety of extracellular particles (EPs), including extracellular vesicles (EVs) and non-vesicular extracellular particles (NVEPs), that transfer a broad range of biomolecules from donor to recipient cells.^1^ Detailed characterization of the biogenesis, release, and uptake of EVs – membranous EPs generated by plasma membrane budding or from intraluminal vesicles of the endosome^2^ – has already demonstrated proof-of-concept for EP-mediated delivery of therapeutic RNAs. For example, surface-engineered exosomes loaded with siRNA repress BACE1 in brain cells, miR-140-loaded exosomes slow the progression of osteoarthritis by targeting chondrocytes, and microvesicles carrying TGF-β1 siRNA increase apoptosis and reduce migration in sarcoma cells.^3–5^

The recent discovery of exomeres, a non-vesicular subclass of EPs, revealed yet another specialized cell messenger implicated in intercellular transfer of RNA and other biomolecules.^6–8^ Exomeres are ∼35-50 nm protein-rich complexes that are naturally enriched in miRNAs and RNA interference (RNAi)-related proteins.^6,7,9^ Two unique features of exomeres suggest that they have considerable potential for use in RNAi therapeutics. First, AGO2 preferentially associates with exomeres as compared to EVs.^6,7^ In human plasma, the majority of miRNAs purify with non-vesicular AGO2 complexes.^10^ Second, miRNAs occupy a substantially higher proportion of the total RNA content in exomeres versus other EP types.^7^ These observations suggest that i) the primary functional role of exomeres is to activate RNAi in recipient cells and ii) in the absence of a protective lipid bilayer, exomeres are sufficiently stable to deliver intact miRNAs to recipient cells. Indeed, it has already been shown that trophoblast-derived exomeres deliver miR-517a-3p to Jurkat cells more effectively than EVs.^8^ Together, these attributes make it important to investigate exomeres as a potential delivery platform for the next-generation of RNAi-based therapeutics.

Careful study of exomeres requires reproducible isolation of a sufficiently pure particle population at reasonable scale. At present, separation of exomeres from other EPs remains a substantial challenge. Asymmetric flow field-flow fractionation (AF4), the technique used to initially discover exomeres, can isolate relatively pure populations but requires fine-tuning for each application.^9,11^ An alternative protocol using differential ultracentrifugation (UC) was introduced in 2023, but it is time-intensive and cannot readily be scaled up to meet the demands of an industrial process.^12^ Moreover, UC can exert significant shear forces on particles, causing fragmentation and making it difficult to determine whether the recovered material accurately reflects what was originally produced by cells.^13^ Most recently, an approach combining fast protein liquid chromatography (FPLC) and size-exclusion chromatography (SEC) was developed to address some of the limitations of UC, underscoring both the importance of studying exomeres and the pressing need for improved isolation methods.^14^ To allow for flexibility in bioprocess development, there remains a significant need for straightforward, scalable exomere isolation processes that yield highly pure populations with minimal processing stress.

Tangential flow filtration (TFF) is a form of ultrafiltration that separates particles based on their hydrodynamic characteristics. TFF systems are relatively inexpensive, allow for successive isolation of different particle types, and can be cleaned for reuse.^15^ TFF has previously been used to isolate a variety of biological species, including EVs, plasmid DNA nanoparticles, and viruses from various biofluids.^16–19^ Here, we developed a tandem TFF method to isolate EVs and exomeres from the conditioned media of HEK293T cells (**Fig. 1**). The objectives of our study were to: (1) develop a tandem TFF protocol for exomere isolation, (2) compare the purity and yield of TFF-isolated particles to particles isolated by ultracentrifugation, and (3) characterize the physical and molecular properties of TFF-isolated exomeres.

**Figure 1:**
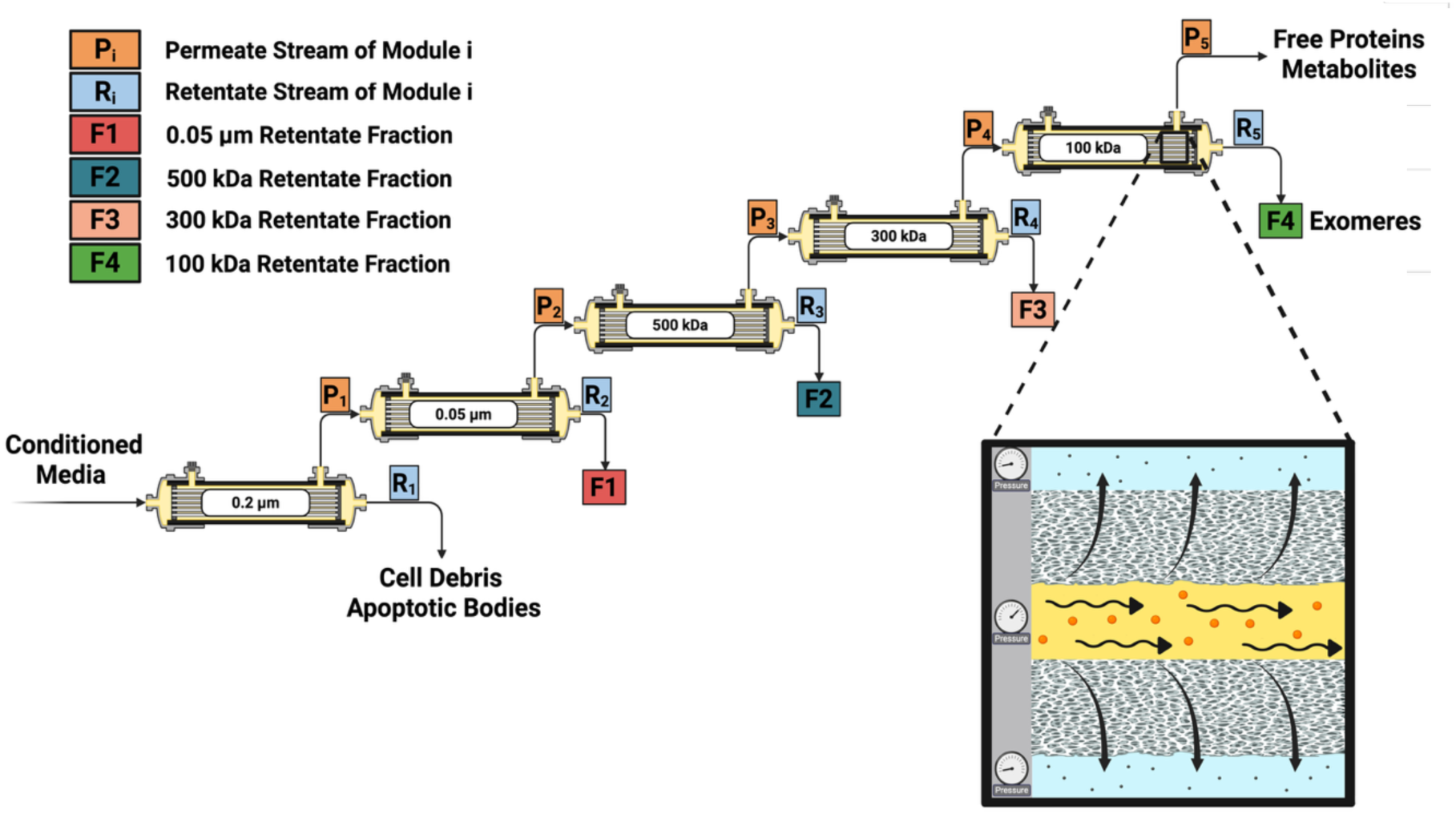
Tandem TFF for isolating exomeres and other extracellular particles from conditioned media. Particle separation is driven by a pressure differential across a semiporous membrane. Conditioned media is first fed to a 0.2 μm molecular weight cutoff (MWCO) clarifying filter to remove cell debris and apoptotic bodies. The permeate is collected and delivered to the next filter in the series. Retained particles on membranes F1-F4 are diafiltered with HEPES-supplemented PBS before collection. Figure created with BioRender.com.

## Experimental Methods

### Cell culture

HEK293T cells (ATCC, CRL-3216) were seeded in high glucose DMEM supplemented with 10% (v/v) FBS and 100 units/mL Penicillin-Streptomycin at a density of 10,000 cells/cm^2^ in 10 cm dishes (small scale) or 18,500 cells/cm^2^ in T225 vented flasks (large scale, optimized conditions) and incubated at 37°C, 5% CO_2_ until the cells were approximately 50-60% confluent. Cells were washed with PBS three times then changed to serum-free DMEM (Thermo, 31053028) supplemented with 100 units/mL Penicillin-Streptomycin, 4 mM L-Glutamine, and 1X Insulin-Transferrin-Selenium (ITS) (Corning, 25-800-CR) for conditioned media production. Conditioned media was collected when cells reached 80-90% confluence and processed with TFF the same day.

#### Whole cell lysate isolation

Cells were trypsinized and pelleted at 300 x g for three minutes. After removal of the supernatant, cells were resuspended in 1 mL of PBS followed by centrifugation at 300 x g for an additional three minutes. The supernatant was removed followed by pellet resuspension in 1 mL of ice-cold RIPA buffer supplemented with EDTA-free Halt^TM^ protease inhibitor cocktail (Thermo, 78425). After briefly vortexing, homogenates were placed on ice for 30 minutes followed by centrifugation for 15 minutes at 14,000 x g, 4°C to clear the lysates. Supernatants were collected and stored at 4°C for a maximum of one week or -80°C long-term.

#### EP isolation by tandem TFF (**Fig. 1**)

Five MicroKros (small scale) or MidiKros (large scale) hollow fiber membranes of MWCOs: 0.2 μm, 0.05 μm - F1, 500 kDa - F2, 300 kDa - F3, and 100 kDa - F4 (Spectrum^®^, Repligen) were preconditioned with DI water followed by PBS containing 25 mM HEPES solution (Sigma-Aldrich, H0887), termed PBS-H.^12^ Preconditioning solutions were used at a volume of 2 mL/cm^2^ membrane surface area. Conditioned media was first clarified with a 0.2 μm filter to remove cell debris and apoptotic bodies. The permeate was collected and fed to the remaining four filters sequentially. The retentate of the 0.2 μm filter and permeate of the 100 kDa filter were discarded.

For particle isolation, retained particles in each module (1-4 mL particle suspension depending on scale) were diafiltered with 8-10 diavolumes of 25 mM PBS-H to remove free proteins and exchange buffers. Fluid volumes were reduced to 1 mL for MicroKros filters or 2-3 mL for MidiKros filters before final collection via syringe. Except for those used in RNA protection assays, particle suspensions were immediately supplemented with EDTA-free protease inhibitor cocktail, inverted briefly to mix, and stored at 4°C until use. Samples were stored at -80°C if not used within one week. Each fraction was designated F1-F4, with smaller numbers corresponding to larger MWCOs.

To regenerate TFF membranes, the filter modules were flushed immediately after use with 2 mL/cm^2^ DI water, followed by 15 mL of 0.5 M NaOH, and then stored in 0.1 M NaOH at 4°C until the next use. Modules were replaced in their entirety when protein content for a given fraction significantly changed, or substantial EV contamination of the non-vesicular fractions was noticed in routine assays. Typically, filters were used for five to seven isolations.

### EP isolation by differential ultracentrifugation (UC)

Differential UC was carried out in the manner previously described.^12^ Briefly, conditioned media was centrifuged for 10 minutes at 500 x g, 4°C to remove cell debris, followed by centrifugation of the supernatant for 20 minutes at 2,000 x g, 4°C to remove apoptotic bodies. The resulting supernatant was centrifuged for 40 minutes at 10,000 x g, 4°C to obtain crude large EVs (LEVs) as a pellet. The pellet was resuspended in PBS-H and washed by an additional centrifugation cycle at 10,000 x g, 4°C. The LEV supernatant was clarified using a 0.2 μm filter, then concentrated with a 100 kDa MWCO centrifugal concentrator (Millipore, UFC710008). Small EVs (sEVs) were isolated by ultracentrifugation of the concentrate, diluted to 37 mL with PBS-H, at 167,000 x g for 4 hours, 4°C in a Beckman Coulter SW 32 Ti swinging-bucket rotor. The crude sEV pellet was washed by suspension of the pellet in 37 mL PBS-H followed by centrifugation at 167,000 x g, 4°C for 4 hours. To isolate exomeres, the sEV supernatant was centrifuged at 167,000 x g, 4°C for 16 hours, resuspended in 37 mL of PBS-H, then washed by centrifugation at 167,000 x g, 4°C for an additional 16 hours. Resuspended particles were stored at 4°C for a maximum of one week or at -80°C for longer periods.

#### Protein quantification

EP samples were quantified without prior lysis. Protein content was estimated using the Pierce^TM^ BCA protein assay (Thermo, 23225) following the manufacturer’s instructions for the microplate procedure, with minor modifications. For incubation, the microplate was kept at 37°C, 5% CO_2_ for two hours instead of 30 minutes.

#### SDS-PAGE and Western blot

Samples were mixed with 4X Laemmli buffer (Bio-Rad, 1610747) at a ratio of 3:1 (v/v) with or without 50 mM DTT (Bio-Rad, 1610611) depending on the antigen, then denatured at 95°C for 5 minutes. For SDS-PAGE, 2.5–10 μg of total protein was loaded per lane onto 4-20% precast polyacrylamide gels (Bio-Rad, 4561094) and separated at 200 volts for approximately 35 minutes. Equal amounts of protein were loaded. Proteins were transferred to 0.2 μm PVDF membranes (Bio-Rad, 1704156) using a Trans-Blot Turbo system with the mixed molecular weight protocol (25 V, 1.3 A, 7 minutes). After transfer, blots were blocked with EveryBlot blocking buffer (Bio-Rad, 12010020) for 1 hour at room temperature with agitation, followed by primary antibody incubation for 12-16 hours at 4°C with agitation. Primary antibodies were diluted (1:1000) in fresh blocking buffer.

After incubation, blots were washed five times for five minutes each in TBST, incubated with secondary antibody in blocking buffer for 1 hour with agitation, and finally washed with TBST an additional six times for five minutes each. Chemiluminescent signals were developed with SuperSignal^TM^ West Pico PLUS substrate (Thermo, 34580) for five minutes before imaging. One additional blot was developed with SuperSignal^TM^ West Femto maximum sensitivity substrate (Thermo, 34094) to probe for potentially weak signals. The following primary antibodies were used: rabbit anti-AGO2 (Abcam, ab186733), rabbit anti-CD9 (Abcam, ab236630), rabbit anti-CD63 (Abcam, ab134045), rabbit anti-FLOT1 (Abcam, ab133497), rabbit anti-HSP90AB1 (Abcam, ab32568), and rabbit anti-CANX (Cell Signaling Technology, 2679). The following secondary antibody was used: goat anti-rabbit IgG, HRP-conjugated (1:20000, Thermo, 31460).

#### Total protein staining

After separating proteins with SDS-PAGE, TFF gels were silver-stained with the Pierce^TM^ Silver Stain Kit (Thermo, 24612), and UC gels fluorescently stained using the Revert^TM^ 700 total protein staining kit (Licor, 926-11010), following manufacturer’s instructions. Gels were imaged on a Bio-Rad Chemidoc system using the silver stain program for silver-stained gels, or StarBright^TM^ B700 program for fluorescently stained gels. Silver-stained gels were uniformly processed in ImageJ 1.54g (Java 1.8.0_345) using a combination of brightness/contrast adjustment, bandpass filtering, and the “enhance contrast” function to reduce overstaining and improve band clarity. Image adjustments did not alter relative protein abundance or band presence/absence.

#### Conventional negative stain transmission electron microscopy (TEM)

EP samples were fixed in 2% (v/v) paraformaldehyde (Electron Microscopy Sciences, 15710) for 20 minutes at room temperature then adsorbed onto 200 mesh formvar coated gold grids (Electron Microscopy Sciences, FCF200-AU) by flotation of the grids on 20 μL droplets of fixed sample for 15 minutes. Grids were washed on three 20 μL droplets of TEM-grade PBS followed by two 20 μL droplets of HPLC grade water, blotting with filter paper after each wash. For contrast, grids were stained with 1% uranyl acetate (Fluka, 94260) for five minutes, air-dried, then imaged on a JEOL JEM-1400Flash at 100 kV.

#### Immunogold TEM

EP samples were fixed and adsorbed onto formvar coated gold grids as described above. For blocking, grids were transferred to 20 μL droplets of 1% BSA (w/v) (Aurion, 900.011) diluted in PBS then incubated at room temperature for 30 minutes. Without washing, grids were transferred to 20 μL droplets of either primary antibody (1:100 Abcam, ab186733 or ab133497) diluted in 1% BSA, or 1% BSA only (negative control), then incubated overnight at 4°C in a Petri dish containing a moist wipe. After primary antibody incubation, grids were washed with three 20 μL droplets of 0.1% BSA, blotted in-between each wash, then transferred to 20 μL droplets of diluted 10 nm gold-conjugated goat anti-rabbit secondary antibody (1:20 Aurion, 25109) and incubated at room temperature for one hour. Grids were washed with three 20 μL droplets of 0.1% BSA in PBS, three 20 μL droplets of HPLC grade water, then stained for contrast with 1% uranyl acetate for five minutes.

#### SNA-I lectin-gold TEM

Exomere samples were fixed, adsorbed, and blocked as before. Instead of primary antibody incubation, samples were incubated with 5 nm gold-conjugated SNA-I lectin, diluted 1:4 (v/v) in 1% BSA in PBS (Ey Labs, GP-6802-5), for 30 minutes at room temperature. Grids were washed with three 20 μL droplets of PBS followed by three 20 μL droplets of HPLC-grade water, then stained for contrast with 1% uranyl acetate. A 1 mg/mL BSA-only grid was prepared as a negative control. Highly sialylated bovine fetuin protein (Millipore Sigma, F2379) was prepared as a positive control. On a fourth grid, exomeres were first incubated with 1 mg/mL unlabeled SNA-I (Ey-labs, L-6802-2) in PBS at room temperature for 30 minutes, followed by incubation with gold-conjugated SNA-I. This grid served as a control for binding specificity of SNA-I lectin.

#### Particle size estimation

UC and TFF exomere particle diameters were estimated using the ImageJ 1.54g (Java 1.8.0_345) “analyze > measure” tool on particles from three separate TEM micrographs each. All particles were measured north to south to reduce bias. The measurement scales were set using the pixel width of the scale bars on the original TEM JPG files. All measurements from each set of micrographs were pooled prior to analysis.

#### RNA protection assay

For RNase A only treatment, EP samples were treated with 1 μg/mL RNase A (Thermo, EN0531), briefly vortexed, then incubated at 37°C for 30 minutes. For Proteinase K/RNase A treatment, samples were treated with 1 μg/mL Proteinase K (Qiagen, RP103B), briefly vortexed, then heated at 56°C for 10 minutes. Samples were immediately quenched with 1X EDTA-free protease inhibitor cocktail (Thermo, 78425) for five minutes at room temperature to inactivate Proteinase K, then treated with 1 μg/mL RNase A as before. For Triton X-100/RNase A treatment, Triton X-100 was added to samples at a final ratio of 0.075% (v/v), vortexed for 30 seconds, then immediately treated with 1 μg/mL RNase A. Optimal Triton X-100 conditions to disrupt EV membranes were determined elsewhere.^20^ As a control for cotreatment effects, GAPDH siRNA from the Qubit^TM^ microRNA assay kit (Thermo, Q32880) was evaluated in parallel. Reagents were prepared fresh each day. A concentration of 1 μg/mL each of RNase A and Proteinase K was selected with consideration of the minimum effective concentration needed to completely degrade the GAPDH siRNA cotreatment control in 30 minutes, and the approximate physiological levels of serine proteases and ribonucleases found in human serum.^21,22^

#### Small RNA extraction and sizing

Small RNAs were isolated from TFF EP samples using the miRNeasy micro kit (Qiagen, 217084) standard protocol, with minor modifications. 120-360 μL of particle suspension was used as input instead of EP pellets. Purified small RNAs were eluted in 30 μL of nuclease-free water after allowing the silica columns to soak for two minutes. This modification improved elution volume consistency across samples. RNA size distributions were qualitatively assessed using the Agilent 5200 Fragment Analyzer and small RNA kit (Agilent, DNF-470-0275). Briefly, purified RNA samples were diluted in nuclease-free water to obtain concentrations within the dynamic range of the assay (25 – 2500 pg/μL, microRNA region). Samples and reference ladder were heat denatured in a thermal cycler at 70°C for 10 minutes then immediately cooled to 4°C and held at 4°C until use. 2 μL of each sample or ladder was added to 18 μL of diluent marker, thoroughly mixed, then separated at 8.0 kV for 24 minutes using protocol DNF-470-33 for a 33 cm capillary array.

#### Proteinase K activity assay

2 μg of BSA dissolved in 25 mM PBS-H was treated with either 1 μg/mL Proteinase K or Proteinase K and 1X protease inhibitor, briefly vortexed, then incubated at 56°C for 10 minutes. One untreated sample was processed in parallel as a control. Proteins were denatured and separated using SDS-PAGE as previously described. The gel was stained with QC colloidal Coomassie stain (Bio-Rad, 1610803) and visualized on a Bio-Rad Chemidoc system using the Coomassie Blue program.

#### RNase A activity assay

300 ng of GAPDH siRNA and a custom oligonucleotide (Eurofins Genomics, sequence: 5’ AUAUAGACAUUACUUAGUAAA 3’) were separately diluted in 25 mM PBS-H. Each sample was treated with 1 μg/mL RNase A, briefly vortexed, then incubated at 37°C for 30 minutes. One untreated sample of each RNA was used as a control. After incubation, RNAs were extracted using the miRNeasy micro kit as described above. RNAs were separated using denaturing nucleic acid PAGE on a 10% TBE-Urea precast gel (Bio-Rad, 4566033) then visualized with SYBR^TM^ Gold (1:10000, Thermo, S11494). The stained gel was imaged on a Bio-Rad Chemidoc system using the SYBR^TM^ Gold program.

#### RNA library generation and sequencing

Small RNAs isolated from TFF EPs were cleared of residual contaminants (Zymo Research, R1013) then used as input for library prep. All RNA sequencing was performed at the Michigan State University RTSF Genomics Core. Libraries were prepared using the Lexogen small RNA-seq library preparation kit for Illumina (Lexogen, 052), following manufacturer’s instructions. Completed libraries were quality controlled and quantified using a combination of Biotium AccuGreen^TM^ high sensitivity dsDNA (Biotium, 31066) and Agilent 4200 Tapestation HS DNA1000 (Agilent, 5067) assays. Libraries were normalized by concentration and pooled in equimolar amounts. The pooled library was then cleaned to remove small fragments and adapter dimers using the Lexogen Purification Module protocol (Lexogen, 022.24) with magnetic beads at a 1.3:1 (v/v) bead-to-library ratio. The library pool was sequenced using an Element Biosciences AVITI Cloudbreak Freestyle 150 cycle medium output kit (Element Biosciences, 860-00014). Sequencing was performed in a 1x50 bp single read format. Base calling was done by AVITI OS v3.3.2 followed by demultiplexing and conversion to FastQ format using Element Biosciences bases2fastq v2.1.0.

#### Small RNA sequencing analysis

Analyses were performed by the Michigan State University Bioinformatics Core. 3’ adapters were removed from all reads using cutadapt.^23^ Untrimmed reads and reads with a length less than 15 bp after trimming were discarded. Reads from each sample were mapped to known miRNAs of the human genome using miRBase to obtain read counts for each miRNA.^24^ This step was carried out using the miRDeep2 pipeline.^25^ The DESeq2 package in R was used to find differentially expressed miRNAs with exomeres taken as the baseline group for log_2_ fold change (LFC), i.e., log_2_(F2 EV mean count/F4 exomere mean count).^26^ The lfcShrink() function was called to shrink the effect size of LFC. The Wald test was used to calculate p-values. Adjusted p-values were calculated using the Benjamini-Hochberg (BH) procedure for FDR control. Differentially expressed miRNAs were taken to be those with an adjusted p-value < 0.05. The counts were normalized using the median of ratios method to account for variations in library size/sequencing depth as well as sample composition.

#### Reverse transcription quantitative real-time PCR (RT-qPCR)

RNA extracted from TFF EP samples using the miRNeasy micro kit (Qiagen, 217084) was cleared of contaminants (Zymo Research, R1013) then used as input. The following TaqMan^TM^ advanced miRNA assays (Thermo, A25576) were used: 478735_mir (hsa-miR-1908-5p), 483118_mir (hsa-miR-12136), 477992_mir (hsa-miR-24-3p), and 477982_mir (hsa-miR-222-3p). Briefly, RNA was 3’ polyadenylated and 5’ adapter ligated then reverse transcribed to generate cDNA using the TaqMan^TM^ Advanced miRNA cDNA synthesis kit (Thermo, A28007). After cDNA amplification using the miR-Amp reaction, qPCR was run on a CFX96 (Bio-Rad). Data were analyzed using the ΔΔC_t_ method with normalization to the geometric mean of the C_t_ values for hsa-miR-24-3p and hsa-miR-222-3p in each sample. F2 EVs were taken as the basis.

#### Statistics

To determine significant differences in TFF fraction total protein abundance, a Kruskal-Wallis nonparametric test with Dunn’s multiple comparisons at α=0.05 was run using R 4.2.2 in BioRender. Sample group means (n=8 per group) for F2, F3, and F4 were compared to F1.

## Results

Recognizing the potential applications of exomeres, we wanted to determine if TFF, which is more compatible with industrial scale bioprocesses than UC and AF4, could reproducibly isolate exomeres from conditioned media. Using the typical size ranges of EVs and exomeres as a guide, we designed a tandem TFF process to separate exomeres from other EPs (**Fig. 1**). We then performed biophysical and molecular characterization on particles isolated from multiple steps in the purification process and benchmarked them against EPs obtained using differential UC.

### Particle retention modeling informs TFF process design

Membrane MWCOs were selected based on particle retention models that assumed a log-normal distribution of pores. Log-normal probability correlations effectively predict the sieving behaviors of synthetic ultrafiltration membranes.^27^ For a 35 nm particle, assumed to be spherical and isotropic, rejection coefficients of 90%, 49%, and 28% were calculated for 5, 15, and 25 nm mean pore size filters, respectively (**Supp. Fig. 1**). We subsequently hypothesized that exomeres could be selectively enriched between a 5 nm and 15 nm filter (designated as retentate fraction F4). For the 15 nm membrane (300 kDa - F3), the calculated rejection coefficients for small (∼60 nm) and large (∼100 nm) EVs were 72% and 88%, respectively. This membrane was subsequently used immediately prior to the terminal filter. Three additional filters sized at 185, 50, and 25 nm (0.2 μm, 0.05 μm - F1, 500 kDa - F2), were used to clarify the conditioned media and isolate EVs upstream of exomeres.

### TFF exomeres are highly pure and enriched in RNAi proteins

Exomere enrichment in fraction F4 was first confirmed using Western blot. F4 was negative for EV markers associated with endocytosis and vesicle trafficking (CD9, CD63, FLOT1) but enriched in RNAi-associated proteins (AGO2, HSP90AB1) (**Fig. 2A**). HSP90 has been proposed as a potential biomarker for mammalian exomeres.^9^ HSP90 helps load small RNA duplexes onto AGO2 in an

**Figure 2:**
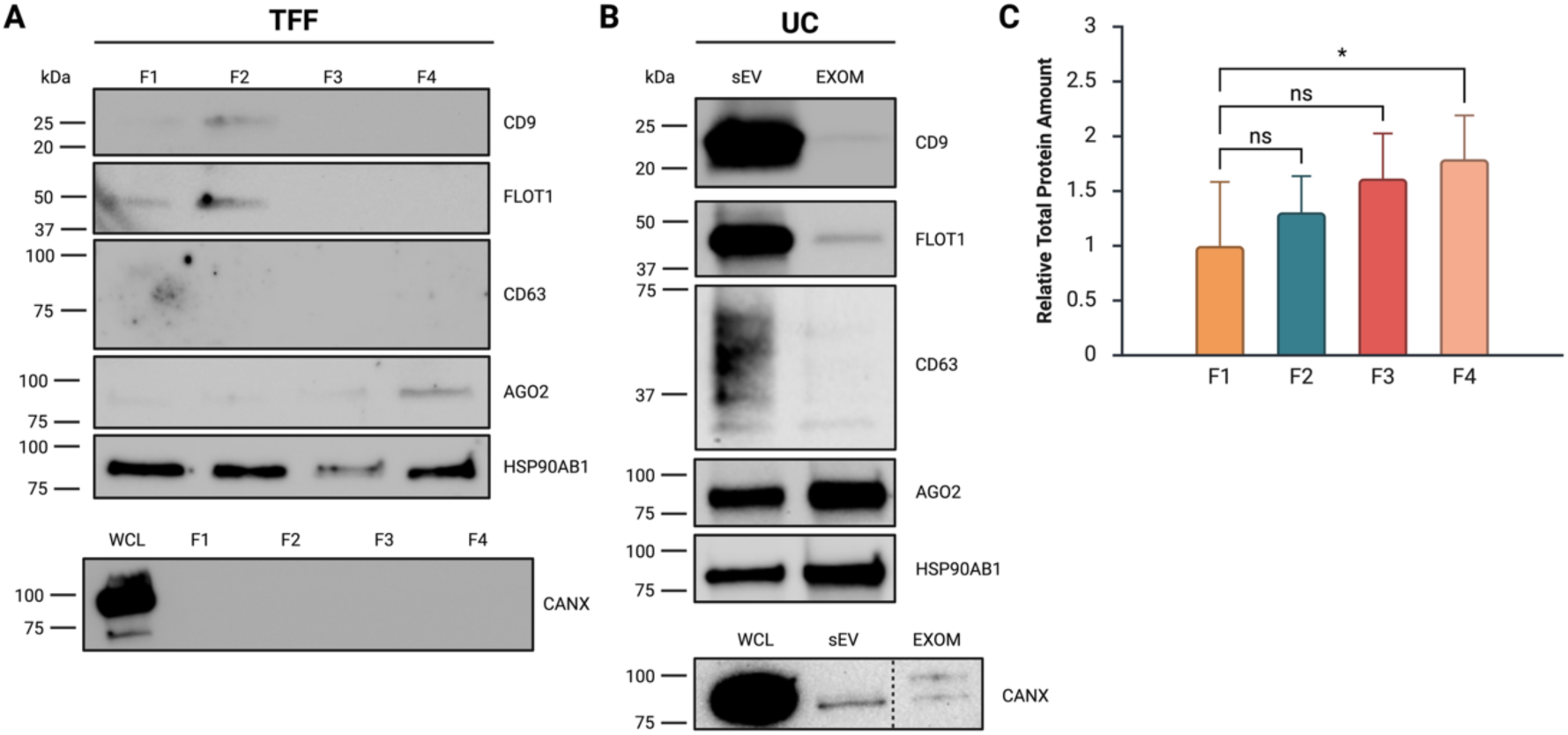
TFF F4 exomeres are comparable in purity and molecular content to UC exomeres. **A.** Western blots of TFF EVs (F1, F2) and non-vesicular particle fractions (F3, F4). Representative blot of three biological replicates shown. **B.** Western blots of differential UC small EVs (sEV) and exomeres (EXOM). Representative blot of two biological replicates shown. The endoplasmic reticulum marker Calnexin (CANX) was tested once in each case as a negative control for co-purification of cell debris. Blots were cropped and uniformly processed in ImageJ 1.54g (Java 1.8.0_345). Unprocessed blots can be found in **Supp. Fig. 2**. **C.** Relative total protein content of F1-F4. Data are presented as mean ± SEM and represent eight biological replicates. A Kruskal-Wallis nonparametric test with Dunn’s multiple comparisons at α=0.05 was run to determine significant differences, * P < 0.05. Figure created with BioRender.com.

ATP-dependent manner and also aids in its targeting to processing bodies and stress granules.^28,29^ F3, containing a less concentrated population of non-membranous nanoparticles, showed faint signals for AGO2 and HSP90 and lacked EV markers. Comparable trends in the depletion and enrichment of protein markers were observed in small EV (sEV) and exomere (EXOM) fractions isolated by UC (**Fig. 2B**). F4 had significantly higher protein content than other fractions, especially compared to large EVs in F1, suggesting that F4 contained protein-rich, non-vesicular particles (**Fig. 2C**).

Total protein staining revealed comparable enrichment of proteins near 15 kDa and 100-150 kDa in both TFF and UC exomere fractions (**Supp. Fig. 2**). Previously, E-PHA lectin blotting detected a high molecular weight glycoprotein near 150 kDa in AsPC-1 and MDA-MB-4175 exomeres.^9^ TFF appeared to yield purer exomeres than UC, as CD9, FLOT1, and CANX (EV and cell debris markers) were consistently detectable at low levels in the UC fractions. We probed fractions F2-F4 on an additional blot with a maximum sensitivity, low-femtogram ECL substrate to confirm that the lack of EV proteins was a function of isolate purity and not inadequate protein loading. We found F3 and F4 to be absent of EV markers in this experiment as well (**Supp. Fig. 2**).

### AGO2 and α2,6-linked sialic acid localize to crenated structures within the TFF exomere fraction

TEM of EVs and exomeres revealed distinct particle morphologies. Both UC and TFF EVs exhibited the typical cup-like structure.^6^ TFF appeared to yield more structurally intact EVs with less aggregation. TFF and UC exomere fractions contained particles of comparable size and morphology. Particle sizes from the UC exomere fraction (49.8 ± 22.0 nm, SD) and TFF exomere fraction (45.6 ± 10.7 nm, SD) were determined by manual measurement of particle diameters from several TEM micrographs (**Supp. Fig. 3**). We attempted to measure particle size using a Particle Metrix ZetaView^®^ nanoparticle tracking analyzer (NTA) but could not detect particles smaller than 50 nm in either UC or TFF exomere fractions, likely due to equipment limitations regarding the detection of very small particles with low refractive indices. TFF produced exomeres with less apparent contamination, as evidenced by the absence of small amorphous aggregates frequently observed in the UC micrographs (**Fig. 3A** and **3B**).

**Figure 3:**
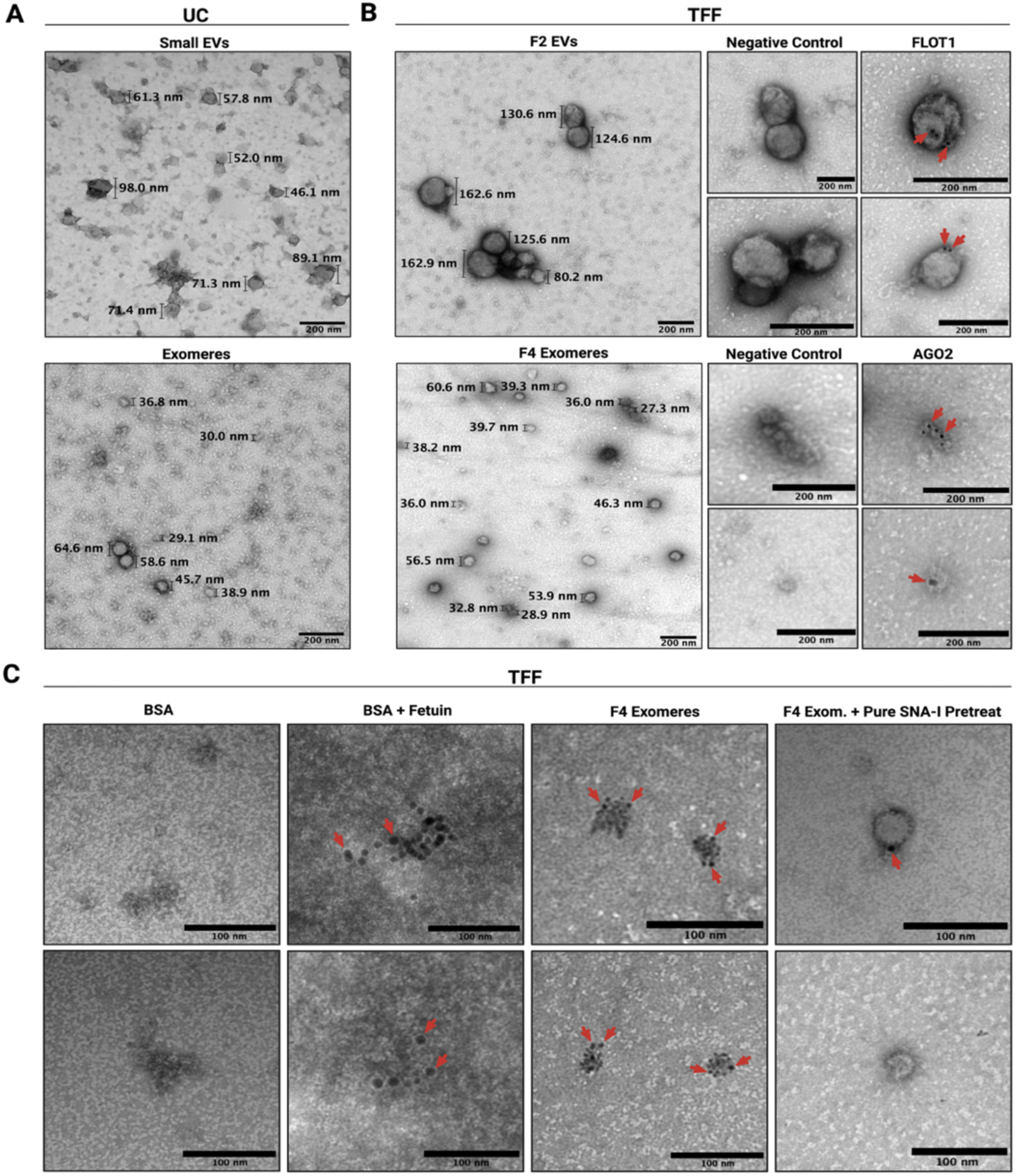
TFF isolates exomeres morphologically similar to those isolated by UC. **A.** Negative-stain TEM of UC sEV and exomere fractions. **B.** Immunogold TEM of TFF EV (F2) and exomere (F4) fractions. **C.** SNA-I lectin gold staining of TFF exomeres. BSA, BSA and fetuin, and pure SNA-I pretreatment were used as controls. Immunogold and lectin-gold experiments were performed twice on independent TFF fractions. Uncropped micrographs are provided in **Supp. Fig. 4**. Figure created with BioRender.com.

Immunogold staining with FLOT1 confirmed the presence of EVs in TFF F2. Treatment of F4 with gold-conjugated AGO2 resulted in the staining of 35-40 nm particles with punctate centers and crenated edges. These structures occurred as both singlets and aggregates (**Fig. 3B**). Similar structures were observed in our UC fraction as well as in prior work on DiFi exomeres.^6,14^ To further investigate if particles stained by AGO2 were exomeres, we performed lectin-gold staining with gold-conjugated SNA-I targeting terminally linked α2,6 sialic acid. Exomeres are major carriers of sialylated glycoproteins.^9^ Lectin-gold particles localized to the same punctate structures as in the AGO2 staining. We confirmed the specificity of SNA-I for α2,6-linked sialic acid using sia-free BSA and a highly-sialylated bovine serum protein, fetuin (**Fig. 3C**). As expected, SNA-I bound to aggregates of fetuin but showed no binding in the BSA-only sample. Pretreatment with unlabeled SNA-I virtually abolished gold labeling of exomeres, indicating that sialic acid sites were already occupied by unconjugated lectin and therefore inaccessible to the conjugate (**Fig. 3C**). The sialoglycoprotein Galectin-3-binding protein (LGALS3BP) is enriched in exomeres and natively self-assembles into ring-like decamers 30-40 nm in size.^9,30^ The structures identified in our lectin-gold experiment were of similar size and strongly bound SNA-I at their perimeters. This suggests that heavily glycosylated proteins might support cell surface-exomere interactions.^9^ These crenated structures were also present in F2 at moderate abundance, indicating that exomeres can copurify with EVs (**Supp. Fig. 3**), as would be expected.

### TFF exomeres protect their endogenous RNAs from nuclease degradation

We next profiled the small RNA content of TFF EVs and exomeres. RNAs in the size range of miRNAs were abundant in both fractions. Larger RNAs (80 – 200 nt) were more prevalent in EVs.

We investigated RNA packaging using an RNase protection assay, in which fractions were treated with combinations of RNase A, Proteinase K, and Triton X-100. Exomere RNAs were remarkably resistant to enzymatic degradation, even in the presence of surfactant or protease. As expected, EV RNAs were substantially degraded upon treatment with Triton X-100, a known EV membrane disruptor (**Fig. 4A** and **4B**).^20^ These data reinforce that exomeres lack membranes but also reveal that exomeres package small RNAs in a highly stable, nuclease-resistant complex. The effects of Triton X-100 and Proteinase K on the activity of RNase A were assessed using a 21-mer GAPDH siRNA. Significant RNA degradation was observed under all treatment conditions, indicating that loss of RNase A activity due to cotreatment was negligible (**Fig. 4C)**.

**Figure 4:**
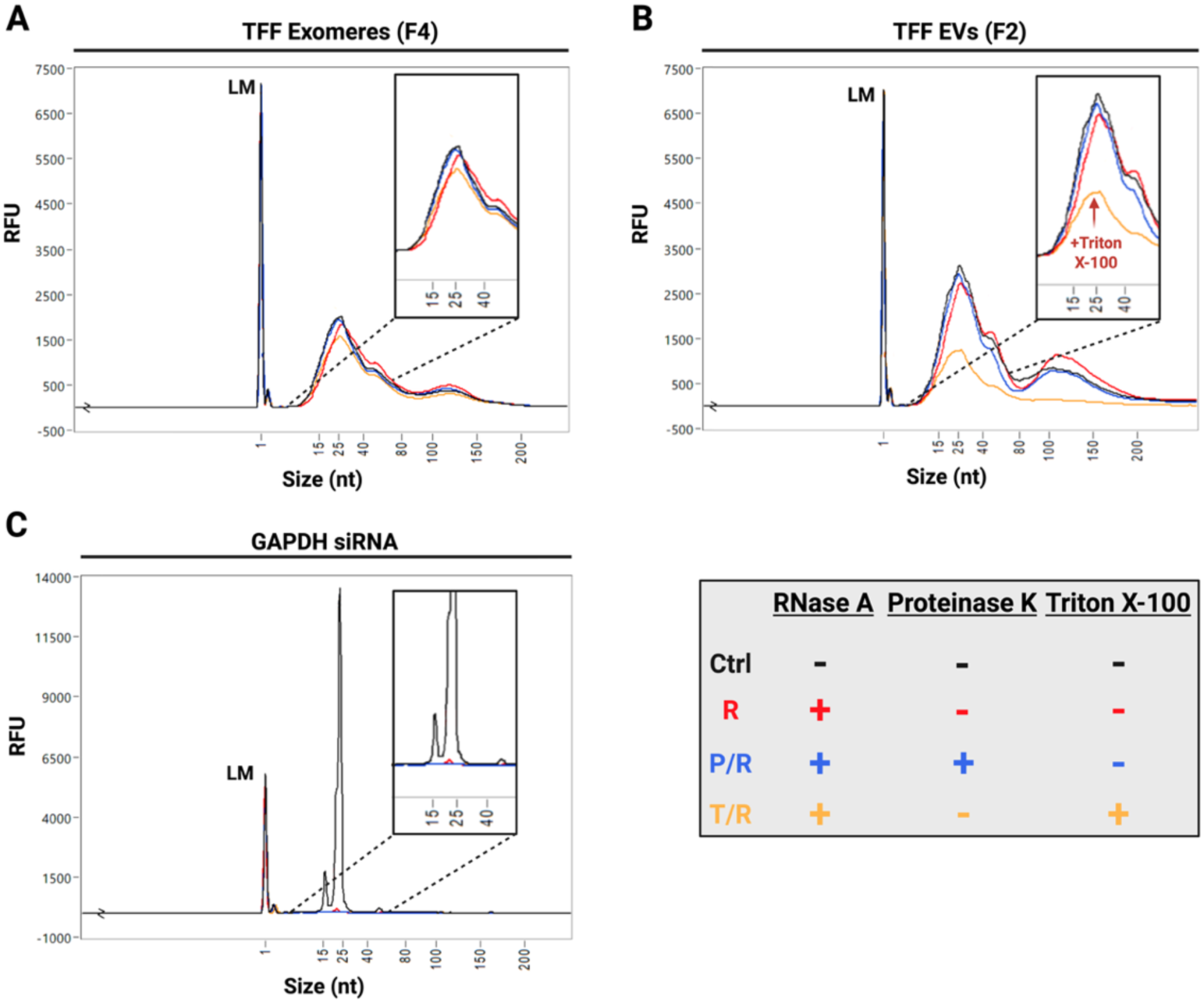
TFF exomere RNAs are protected against enzymatic degradation in the presence of protease and surfactant. Representative fragment analyzer traces of small RNAs isolated from **A.** TFF exomeres (F4) and **B.** TFF EVs (F2). Treatment conditions: Ctrl (Control), R (RNase A only), P/R (Proteinase K/RNase A), T/R (Triton X-100/RNase A). **C.** GAPDH siRNA measured in parallel as a control. Replicate experiments can be found in **Supp. Fig. 5**. Figure created with BioRender.com.

Proteinase K activity was verified by treating BSA with either Proteinase K, or Proteinase K and protease inhibitor, followed by SDS-PAGE and Coomassie blue staining. As expected, Proteinase K degraded BSA, with degradation being limited by the presence of protease inhibitor (**Supp. Fig. 5**). We also confirmed that RNase A could cleave single- and double-stranded RNAs in our TFF buffer (25 mM PBS-H), using GAPDH siRNA for the double-strand and a scrambled RNA oligomer designed to avoid hairpin formation and palindromic basepairing for the single-strand. RNase A degraded both RNAs but showed a stronger activity against the single-stranded RNA, further indicating that exomere RNAs, which are likely single-stranded, are protected by a nuclease-resistant complex (**Supp. Fig. 5**).

### TFF EVs and exomeres are enriched in distinct populations of miRNAs

To confirm the presence of miRNAs in our TFF EV and exomere samples, we performed RNA sequencing on the F2 and F4 small RNA fractions. Overall, sequencing reads for EVs and exomeres were successfully mapped to 812 mature miRNAs using the miRBase microRNA database.^24^ 27 miRNAs were differentially expressed, six of which were significantly more enriched in exomeres (**Fig. 5A**). miR-4516 and miR-3960 were the most abundant miRNAs in each fraction by a considerable number of reads. miR-320a-3p, a highly abundant miRNA in DiFi exomeres and supermeres^7^, was also abundant in our TFF exomeres as well as differentially expressed compared to TFF EVs (**Fig. 5A** and **5B**). Surprisingly, miR-30a-3p was found only in the exomere fraction across all replicates.

**Figure 5:**
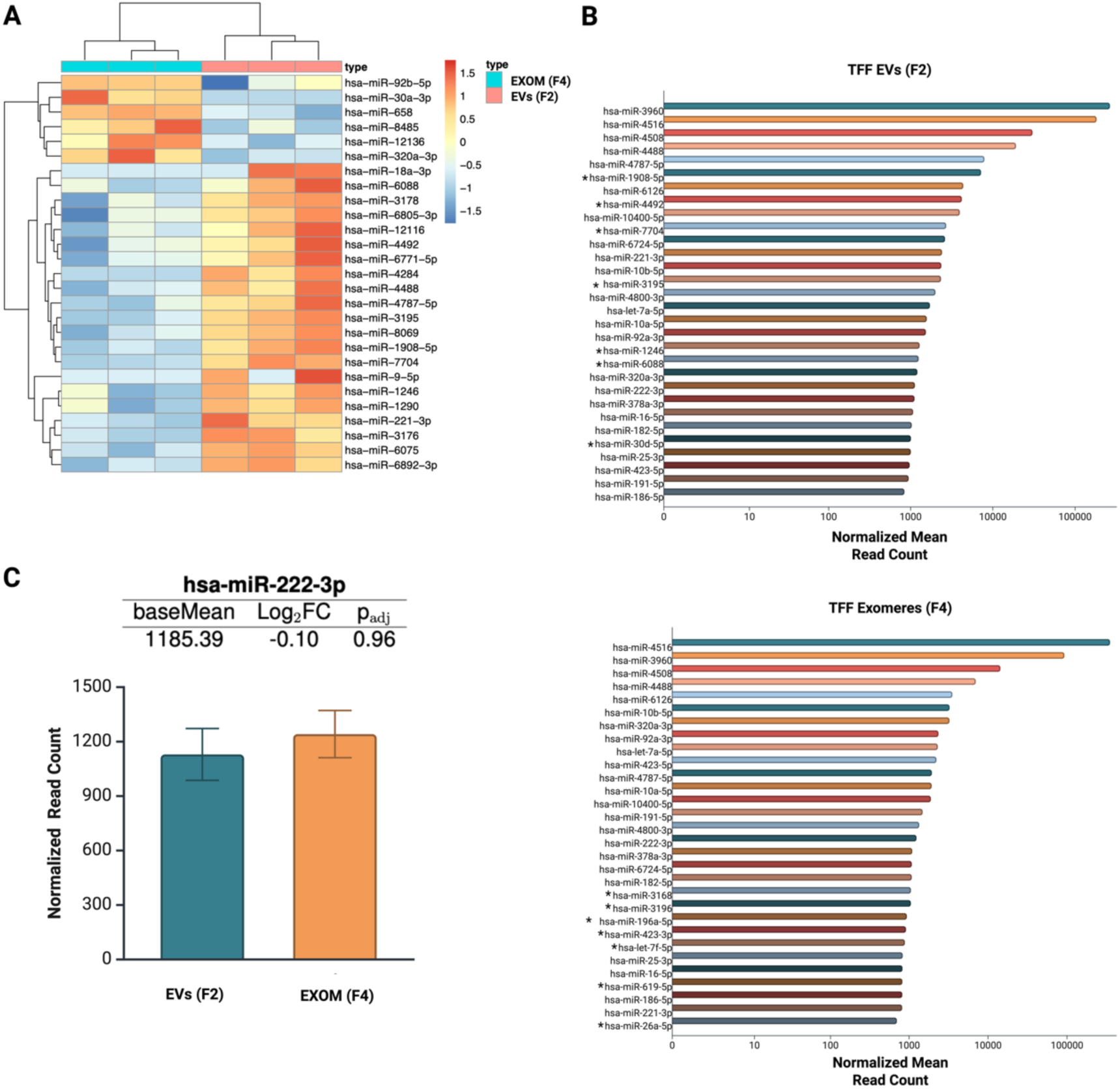
TFF EVs and exomeres are enriched in distinct populations of microRNAs. **A.** Differentially expressed miRNAs between EVs and exomeres, p_adj_ < 0.05. Data represent three biological replicates. **B.** 30 most abundant miRNAs in EVs (top) and exomeres (bottom) based on normalized mean read counts. Asterisks (*) indicate miRNAs unique to EVs (top) or exomeres (bottom) within the 30 most abundant. **C.** Differential expression of hsa-miR-222-3p. Data are presented as mean ± SD and represent three biological replicates. Data were normalized using the median of ratios method. Figure created with BioRender.com.

Stable reference molecules (e.g., proteins or nucleic acids) suitable for normalization across EP types have yet to be identified. Accordingly, we mined our RNA-seq data to identify potential endogenous controls for use in routine assays. miR-222-3p and miR-24-3p, abundant and stably expressed miRNAs found in DiFi cells, sEVs, exomeres, and supermeres^7^, were identified as potential candidates. miR-222-3p ranked among the top 30 most abundant miRNAs in both TFF EVs and exomeres and showed comparable abundance between samples based on normalized read counts and fold change (**Fig. 5C**). This observation was further confirmed using TaqMan^TM^ probe RT-qPCR, indicating that both miR-222-3p and miR-24-3p may be appropriate reference molecules for normalizing changes in miRNA expression across different EP types moving forward. It is a best practice to use multiple internal controls when normalizing qPCR data.^31^ We verified that differential expression trends were consistent between RNA-seq and RT-qPCR using miR-1908-5p and miR-12136 (**Supp. Fig. 6**). Selection criteria for targets to validate were: fold change, FDR adjusted p-value, and a normalized mean read count of at least 100 reads for each sample group. One hundred reads per group has been suggested as a suitable threshold for selecting miRNA candidates for qPCR validation.^32^ Overall, our findings reveal that exomeres isolated by tandem TFF are biologically comparable to exomeres isolated by UC (by us and others), of greater purity than those isolated by UC, and contain small RNAs that are resistant to RNase degradation even in the presence of surfactant and protease.

## Discussion

Ultracentrifugation is currently the most well-described approach for isolating exomeres from conditioned media and is more accessible than AF4. However, UC is time-intensive, risks exomere co-sedimentation with EVs and other components, and cannot easily be scaled to produce pilot or production scale quantities of exomeres. Recently, fast protein liquid chromatography (FPLC)-based size exclusion chromatography (SEC) was introduced as a more scalable alternative to UC and AF4 for the isolation of exomeres and supermeres, underscoring the importance of studying exomeres and the urgent need for improved isolation methods.^14^ To provide another rapid and scalable method for the isolation of mammalian exomeres, we developed a tandem TFF process that yields highly pure exomeres within hours of media harvest. 360 mL of conditioned media was routinely processed in as little as three hours. Indeed, other efforts to apply TFF to isolate NVEPs are emerging^33^, reinforcing the potential of our purification approach.

TFF exomeres displayed molecular and morphological features consistent with exomeres isolated by UC, including the enrichment of RNAi proteins AGO2 and HSP90AB1. The absence of FLOT1, CD9, and CANX signals on Western blots, as well as reduced TEM background, indicated that TFF produced fractions of higher purity relative to UC. LGALS3BP, a glycosylated protein with a high concentration of terminally linked sialic acid, is known to be enriched in exomeres.^9^ SNA-I lectin-gold staining targeting α2,6-linked sialic acid resulted in the localization of gold particles to the exomere perimeter, suggesting a non-random distribution of sialic acid on the particle surface that may serve to mediate interactions with recipient cells. These observations may further suggest that the indented core of the exomere is a structural feature rather than a technical artifact. LGALS3BP self-assembles into 30-40 nm ring-like oligomers, possibly enabling the encapsulation of protein complexes.^30^ Given the localization of gold-conjugated AGO2 to the same types of crenated structures stained by SNA-I, it is tempting to speculate that glycoprotein-AGO2 complexes might be a fundamental constituent of the exomere. Future studies will investigate whether oligomers of glycosylated proteins copurify with AGO2 and other notable components of exomeres such as miRNAs.

Exomeres naturally transport components of RNAi, making them promising candidates for RNAi therapeutics and biomarker discovery (e.g., of differentially incorporated miRNAs). Because miRNAs must persist in circulation to be functional in target tissues, we attempted to degrade the small RNAs in TFF EVs and exomeres using RNase A combined with either the membrane disruptor Triton X-100 or the broad-spectrum protease, Proteinase K. Remarkably, exomere RNAs persisted through each treatment. Conversely, EV RNAs were substantially degraded by RNase A after Triton X-100 treatment, reinforcing that exomeres and EVs are distinct in their structure and therefore may differ in their endocytosis and processing by recipient cells.

RNA sequencing confirmed that miRNAs were abundant in both particle types. A total of 812 mature miRNAs were successfully mapped across EV and exomere sequence reads, six of which were preferentially associated with exomeres. Surprisingly, miR-30a-3p was found only in the exomere fractions across all replicates. miR-30a-3p has been shown to suppress the invasion potential of bladder cancer cells and improve their chemosensitivity *in vitro*, suggesting a potential therapeutic role for exomeres in cancer treatment.^34^ It is worth noting that the most abundant miRNAs in each fraction were those with exceptionally high GC content (>85%, top 4 miRNAs ranked by normalized mean read count). High GC content in the canonical miRNA seed region (i.e., positions 2-8 from the 5’ end) and the extended seed region (positions 4-10) strongly correlates with perfect base pairing to the mRNA target, and an increase in the number of matched seed pairs improves the thermodynamic stability of the miRNA-mRNA duplex.^35^ Given sufficient time for RISC to recruit deadenylation machinery, a translationally repressed transcript can further be destabilized and degraded, eliminating the possibility of its recovery via cytoplasmic polyadenylation.^36^ For intercellular miRNAs that are i) in low adundance relative to intracellular levels and ii) susceptible to degradation by circulating nucleases, high GC content may be evolutionarily favored for enhancing potency and increasing the likelihood of complete target inactivation. This mechanism would be consistent with the known functions of EPs, as cells must communicate clear and active signals to neigboring cells to rapidly enact change in response to stress or other stimuli. To validate a subset of our sequencing results with RT-qPCR, we tested two differentially expressed miRNAs, miR-1908-5p and miR-12136, as well as two that were stably expressed, miR-24-3p and miR-222-3p, and found those results to be in general agreement with the RNA sequencing. miR-24-3p and miR-222-3p appear to be useful endogenous controls for assays that require stable reference RNAs.

Our results further support that exomeres and EVs are produced through overlapping (due to shared molecular content) but distinct biogenesis mechanisms. Previous work demonstrated that sequence motifs direct some miRNAs to EVs, and their loading is mediated by a sumoylated form of the ribonucleoprotein hnRNPA2B1.^37^ However, hnRNPA2B1 appears to preferentially associate with NVEPs.^7^ AGO2 phosphorylation was previously found to regulate the packaging of certain miRNAs into exosomes.^38^ We and others found AGO2 to be mainly associated with the non-vesicular fractions.^6–8^ Further characterization of individual RNA-binding proteins in each EP fraction will be needed to better understand the biogenesis of different classes of EPs, as bulk characterization obscures the link between specific intracellular signaling and sorting processes and the biogenesis of individual extracellular particles.

Moving forward, we are continuing to refine the TFF process for further purification and characterization of exomeres. Although exomeres must be collected in serum-free media, using ITS as a serum substitute complicated biophysical characterization because its protein components persisted through TFF. Transferrin has been shown to form fibrillar deposits on carbon-coated formvar surfaces in the size range of EVs, which may have complicated visual assessments of purity using TEM.^39^ Additionally, we cannot completely rule out the presence of other extracellular particles in our exomere fraction. Lipoprotein complexes are secreted particles of similar size to exomeres. Intermediate-density lipoproteins (IDLs) and very-low-density lipoproteins (VLDLs) appear as 20-60 nm particles under TEM.^40^ However, we are reasonably confident that the particles we isolated were exomeres, since i) HEK293 cells do not express apolipoproteins according to the Human Protein Atlas^41,42^; and ii) TEM grids containing only BSA, the blocking reagent for gold staining experiments, and fresh ITS media were free from particles with the exomere morphology.

We believe this separation process can be applied to other moderately viscous biofluids such as serum and lymph. TFF has already been employed to isolate protein complexes from human plasma and human Fraction IV (FIV), and EVs from lipoaspirate.^16,43^ Apart from RNAi therapeutics, exomeres have potential utility as circulating biomarkers. This TFF process could allow for the rapid isolation and analysis of exomeres from patient biofluids, potentially leading to the discovery of novel disease biomarkers.

## Conclusions

This study proposes a scalable, rapid, and cost-effective method to isolate highly pure EVs and exomeres from conditioned media, enabling further research toward applications in RNAi therapeutics. This study also demonstrates, for the first time, that exomeres protect their constituent miRNAs more comprehensively from nuclease degradation relative to EVs, reinforcing that they likely play a complementary role to EVs in supporting intercellular communication.

## Author contributions

TS contributed to conceptualization, formal analysis, investigation, methodology, writing the original draft, and reviewing/editing of the final work.

YH contributed to conceptualization, funding acquisition, investigation, methodology, and reviewing/editing of the final work.

OB contributed to investigation, validation of results, and reviewing/editing of the final work. CC, MH, and SPW contributed to conceptualization, funding acquisition, project administration, resources, supervision, and reviewing/editing of the final work.

All authors read and approved the final manuscript.

## Declarations

### Data availability

Data generated or analyzed during this study are included in this published article and its supplementary files. RNA sequencing data can be found in the SRA database at BioProject accession: PRJNA1347467.

## Acknowledgments

This work was supported by the National Institutes of Health (Award No. **R21GM154180, 1R01CA286786**); the National Science Foundation (Award No. CBET **2232658**); and the Eunice Kennedy Shriver National Institute of Child Health & Human Development of the National Institutes of Health (Award No. **T32HD087166).** Microscopy data was collected at the MSU Center for Advanced Microscopy. We also acknowledge the genomics services performed in the MSU RTSF Genomics Core as well as the RNA sequencing analyses performed by Dr. Nanye Long at the Michigan State University Bioinformatics Core (RRID:SCR_026706).

## References

1. S. Chen, Q. Bao, W. Xu and X. Zhai, J Nanobiotechnol, 2025, 23, 263.

2. E. R. Abels and X. O. Breakefield, Cell Mol Neurobiol, 2016, 36, 301–312.

3. L. Alvarez-Erviti, Y. Seow, H. Yin, C. Betts, S. Lakhal and M. J. A. Wood, Nat Biotechnol, 2011, 29, 341–345.

4. Y. Liang, X. Xu, X. Li, J. Xiong, B. Li, L. Duan, D. Wang and J. Xia, ACS Appl. Mater. Interfaces, 2020, 12, 36938–36947.

5. Y. Zhang, L. Li, J. Yu, D. Zhu, Y. Zhang, X. Li, H. Gu, C.-Y. Zhang and K. Zen, Biomaterials, 2014, 35, 4390–4400.

6. Q. Zhang, J. N. Higginbotham, D. K. Jeppesen, Y.-P. Yang, W. Li, E. T. McKinley, R. Graves-Deal, J. Ping, C. M. Britain, K. A. Dorsett, C. L. Hartman, D. A. Ford, R. M. Allen, K. C. Vickers, Q. Liu, J. L. Franklin, S. L. Bellis and R. J. Coffey, Cell Reports, 2019, 27, 940–954.e6.

7. Q. Zhang, D. K. Jeppesen, J. N. Higginbotham, R. Graves-Deal, V. Q. Trinh, M. A. Ramirez, Y. Sohn, A. C. Neininger, N. Taneja, E. T. McKinley, H. Niitsu, Z. Cao, R. Evans, S. E. Glass, K. C. Ray, W. H. Fissell, S. Hill, K. L. Rose, W. J. Huh, M. K. Washington, G. D. Ayers, D. T. Burnette, S. Sharma, L. H. Rome, J. L. Franklin, Y. A. Lee, Q. Liu and R. J. Coffey, Nat Cell Biol, 2021, 23, 1240–1254.

8. S. Noguchi, S. Tozawa, T. Sakurai, A. Ohkuchi, H. Takahashi, H. Fujiwara and T. Takizawa, Journal of Reproductive Immunology, 2024, 161, 104187.

9. H. Zhang, D. Freitas, H. S. Kim, K. Fabijanic, Z. Li, H. Chen, M. T. Mark, H. Molina, A. B. Martin, L. Bojmar, J. Fang, S. Rampersaud, A. Hoshino, I. Matei, C. M. Kenific, M. Nakajima, A. P. Mutvei, P. Sansone, W. Buehring, H. Wang, J. P. Jimenez, L. Cohen-Gould, N. Paknejad, M. Brendel, K. Manova-Todorova, A. Magalhães, J. A. Ferreira, H. Osório, A. M. Silva, A. Massey, J. R. Cubillos-Ruiz, G. Galletti, P. Giannakakou, A. M. Cuervo, J. Blenis, R. Schwartz, M. S. Brady, H. Peinado, J. Bromberg, H. Matsui, C. A. Reis and D. Lyden, Nat Cell Biol, 2018, 20, 332–343.

10. J. D. Arroyo, J. R. Chevillet, E. M. Kroh, I. K. Ruf, C. C. Pritchard, D. F. Gibson, P. S. Mitchell, C. F. Bennett, E. L. Pogosova-Agadjanyan, D. L. Stirewalt, J. F. Tait and M. Tewari, Proc. Natl. Acad. Sci. U.S.A., 2011, 108, 5003–5008.

11. E. Willms, C. Cabañas, I. Mäger, M. J. A. Wood and P. Vader, Front. Immunol., 2018, 9, 738.

12. Q. Zhang, D. K. Jeppesen, J. N. Higginbotham, J. L. Franklin and R. J. Coffey, Nat Protoc, 2023, 18, 1462–1487.

13. R. Štukelj, K. Schara, A. Bedina-Zavec, V. Šuštar, M. Pajnič, L. Pađen, J. L. Krek, V. Kralj-Iglič, A. Mrvar-Brečko and R. Janša, European Journal of Pharmaceutical Sciences, 2017, 98, 17–29.

14. O. S. Tutanov, C. Massick, M. Ramirez, J. N. Higginbotham, L. Jimenez, M. Castleberry, Z. Cao, E. John, M. S. Hamilton, Q. Zhang, D. K. Jeppesen, D. L. Michell, S. E. Glass, P. Patel, K. L. Rose, E. Krystofiak, H.-C. Chen, Q. Sheng, Q. Liu, J. G. Patton, A. M. Weaver, J. L. Franklin, K. C. Vickers and R. J. Coffey, Cell Reports, 2025, 44, 116287.

15. P. Agrawal, K. Wilkstein, E. Guinn, M. Mason, C. I. Serrano Martinez and J. Saylae, Org. Process Res. Dev., 2023, 27, 571–591.

16. S. Busatto, G. Vilanilam, T. Ticer, W.-L. Lin, D. Dickson, S. Shapiro, P. Bergese and J. Wolfram, Cells, 2018, 7, 273.

17. Y. Kawai-Harada, V. Nimmagadda and M. Harada, BMC Methods, 2024, 1, 9.

18. H.-W. Liu, Y. Hu, Y. Ren, H. Nam, J. L. Santos, S. Ng, L. Gong, M. Brummet, C. A. Carrington, C. G. Ullman, M. G. Pomper, I. Minn and H.-Q. Mao, ACS Appl. Mater. Interfaces, 2021, 13, 30326–30336.

19. S. Higuchi, T. Satou and Y. Uchida, MethodsX, 2023, 10, 102126.

20. X. Osteikoetxea, B. Sódar, A. Németh, K. Szabó-Taylor, K. Pálóczi, K. V. Vukman, V. Tamási, A. Balogh, Á. Kittel, É. Pállinger and E. I. Buzás, Org. Biomol. Chem., 2015, 13, 9775–9782.

21. J. M. G. Artigas, M. E. Garcia, M. R. A. Faure and A. M. B. Gimeno, Postgraduate Medical Journal, 1981, 57, 219–222.

22. J. C. Houck and L. B. Berman, Journal of Applied Physiology, 1958, 12, 473–476.

23. M. Martin, EMBnet j., 2011, 17, 10.

24. A. Kozomara, M. Birgaoanu and S. Griffiths-Jones, Nucleic Acids Research, 2019, 47, D155–D162.

25. M. R. Friedländer, S. D. Mackowiak, N. Li, W. Chen and N. Rajewsky, Nucleic Acids Research, 2012, 40, 37–52.

26. M. I. Love, W. Huber and S. Anders, Genome Biol, 2014, 15, 550.

27. J. Ren, Z. Li and F.-S. Wong, Journal of Membrane Science, 2006, 279, 558–569.

28. S. Iwasaki, M. Kobayashi, M. Yoda, Y. Sakaguchi, S. Katsuma, T. Suzuki and Y. Tomari, Molecular Cell, 2010, 39, 292–299.

29. J. M. Pare, N. Tahbaz, J. López-Orozco, P. LaPointe, P. Lasko and T. C. Hobman, MBoC, 2009, 20, 3273–3284.

30. S. A. Müller, T. Sasaki, P. Bork, B. Wolpensinger, T. Schulthess, R. Timpl, A. Engel and J. Engel, Journal of Molecular Biology, 1999, 291, 801–813.

31. J. Vandesompele, K. De Preter, F. Pattyn, B. Poppe, N. Van Roy, A. De Paepe and F. Speleman, Genome Biol, 2002, 3, research0034.1.

32. V. Mussack, S. Hermann, D. Buschmann, B. Kirchner and M. W. Pfaffl, in Quantitative Real-Time PCR, eds. R. Biassoni and A. Raso, Springer New York, New York, NY, 2020, vol. 2065, pp. 23–38.

33. M. J. Rogers, A. Andreosso, J. Billakanti, M. G. Duke, Y. Lu, N. Collinson, J. Tan, L. Macia, N. Koifman, M. Floetenmeyer, M. K. Jones, C. A. Gordon and S. Navarro, 2025, preprint, DOI: 10.1101/2025.03.24.645107.

34. T. I.-S. Hwang, P.-C. Chen, T.-F. Tsai, J.-F. Lin, K.-Y. Chou, C.-Y. Ho, H.-E. Chen and A.-C. Chang, Cell Death Dis, 2022, 13, 390.

35. X. Wang, Bioinformatics, 2014, 30, 1377–1383.

36. S. W. Eichhorn, H. Guo, S. E. McGeary, R. A. Rodriguez-Mias, C. Shin, D. Baek, S. Hsu, K. Ghoshal, J. Villén and D. P. Bartel, Molecular Cell, 2014, 56, 104–115.

37. C. Villarroya-Beltri, C. Gutiérrez-Vázquez, F. Sánchez-Cabo, D. Pérez-Hernández, J. Vázquez, N. Martin-Cofreces, D. J. Martinez-Herrera, A. Pascual-Montano, M. Mittelbrunn and F. Sánchez-Madrid, Nat Commun, 2013, 4, 2980.

38. A. J. McKenzie, D. Hoshino, N. H. Hong, D. J. Cha, J. L. Franklin, R. J. Coffey, J. G. Patton and A. M. Weaver, Cell Reports, 2016, 15, 978–987.

39. C. Booyjzsen, C. A. Scarff, B. Moreton, I. Portman, J. H. Scrivens, G. Costantini and P. J. Sadler, Biochimica et Biophysica Acta (BBA) - General Subjects, 2012, 1820, 427–436.

40. L. Zhang, J. Song, G. Cavigiolio, B. Y. Ishida, S. Zhang, J. P. Kane, K. H. Weisgraber, M. N. Oda, K.-A. Rye, H. J. Pownall and G. Ren, Journal of Lipid Research, 2011, 52, 175–184.

41. M. Uhlén, L. Fagerberg, B. M. Hallström, C. Lindskog, P. Oksvold, A. Mardinoglu, Å. Sivertsson, C. Kampf, E. Sjöstedt, A. Asplund, I. Olsson, K. Edlund, E. Lundberg, S. Navani, C. A.-K. Szigyarto, J. Odeberg, D. Djureinovic, J. O. Takanen, S. Hober, T. Alm, P.-H. Edqvist, H. Berling, H. Tegel, J. Mulder, J. Rockberg, P. Nilsson, J. M. Schwenk, M. Hamsten, K. Von Feilitzen, M. Forsberg, L. Persson, F. Johansson, M. Zwahlen, G. Von Heijne, J. Nielsen and F. Pontén, Science, 2015, 347, 1260419.

42. The Human Protein Atlas, https://www.proteinatlas.org, (accessed October 2025).

43. I. S. Pires and A. F. Palmer, Journal of Membrane Science, 2021, 618, 118712.

