## Supplemental Data for "A scalable filtration-based method for isolating exomeres and other nanoscale extracellular particles"

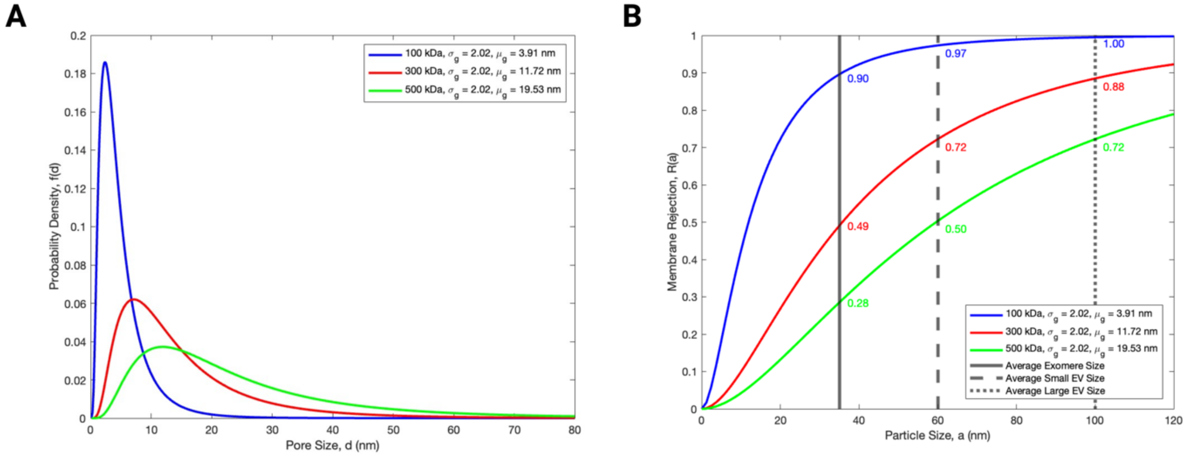


**Supplementary Figure 1:** Filter rejection predictions using log-normal models previously described.[1] **A.** Probability density functions showing the theoretical pore size distribution for 100, 300, and 500 kDa MWCO membranes. **B.** Membrane rejection performance for particles with a size of 35 nm (exomeres), 60 nm (sEVs), and 100 nm (LEVs). The value chosen for geometric standard deviation ($\sigma_g$) was adopted from prior literature based on measurements of asymmetric, modified polyethersulfone (PES) membranes which well-represent the commercial membranes used in this study.[2] Geometric means ($\mu_g$) were calculated from $\sigma_g$ and the mean pore diameter reported by the manufacturer.

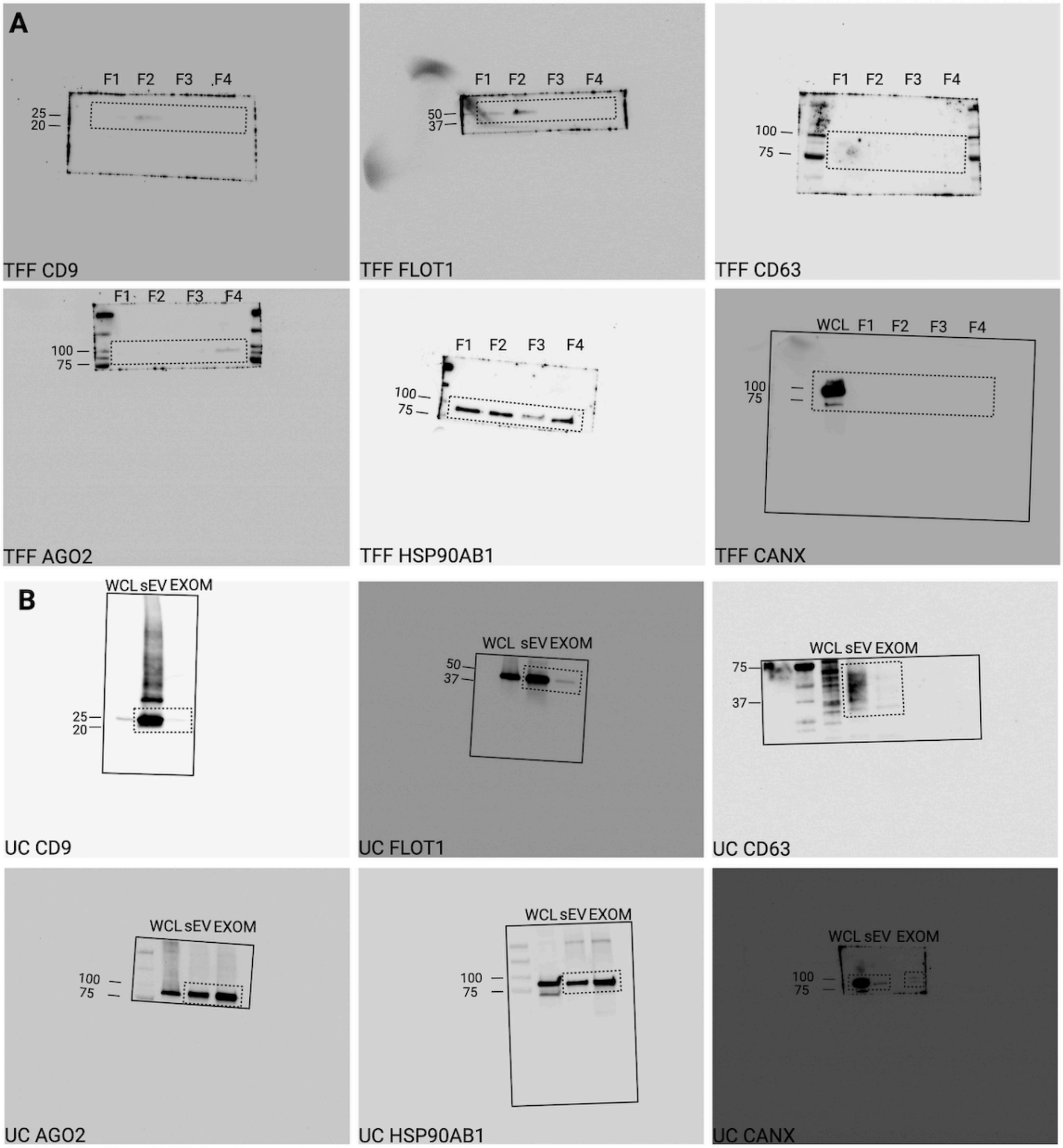


**Supplementary Figure 2:** Unprocessed Western blots for **A.** Tandem TFF and **B.** Differential UC. Dotted boxes represent the regions of the blots presented in **Fig. 2**. Solid boxes represent the outline of the complete blot.

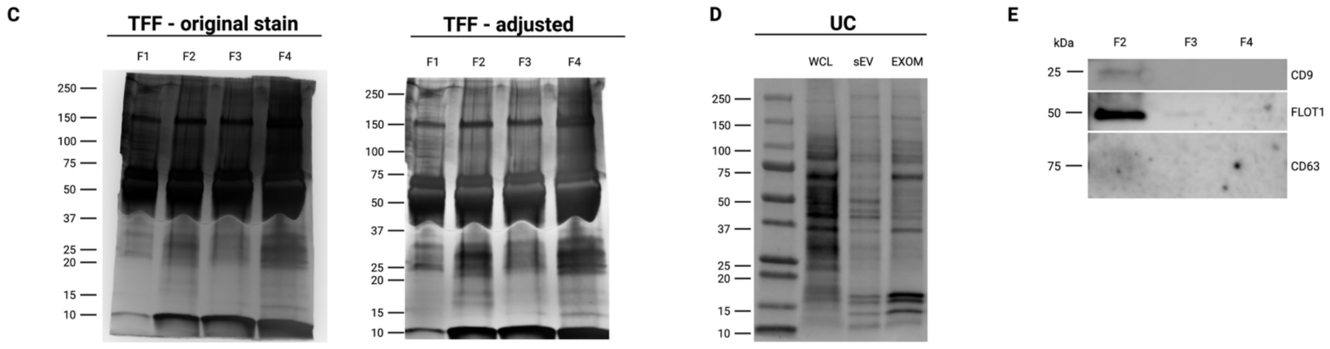


**Supplementary Figure 2 (continued):** Total protein visualization by **C**. silver staining for TFF fractions and **D.** Revert700 fluorescent staining for UC fractions. **E.** Femto ECL Western blot showing absence of EV markers in NVEP fractions F3 and F4 in comparison to EVs in F2.

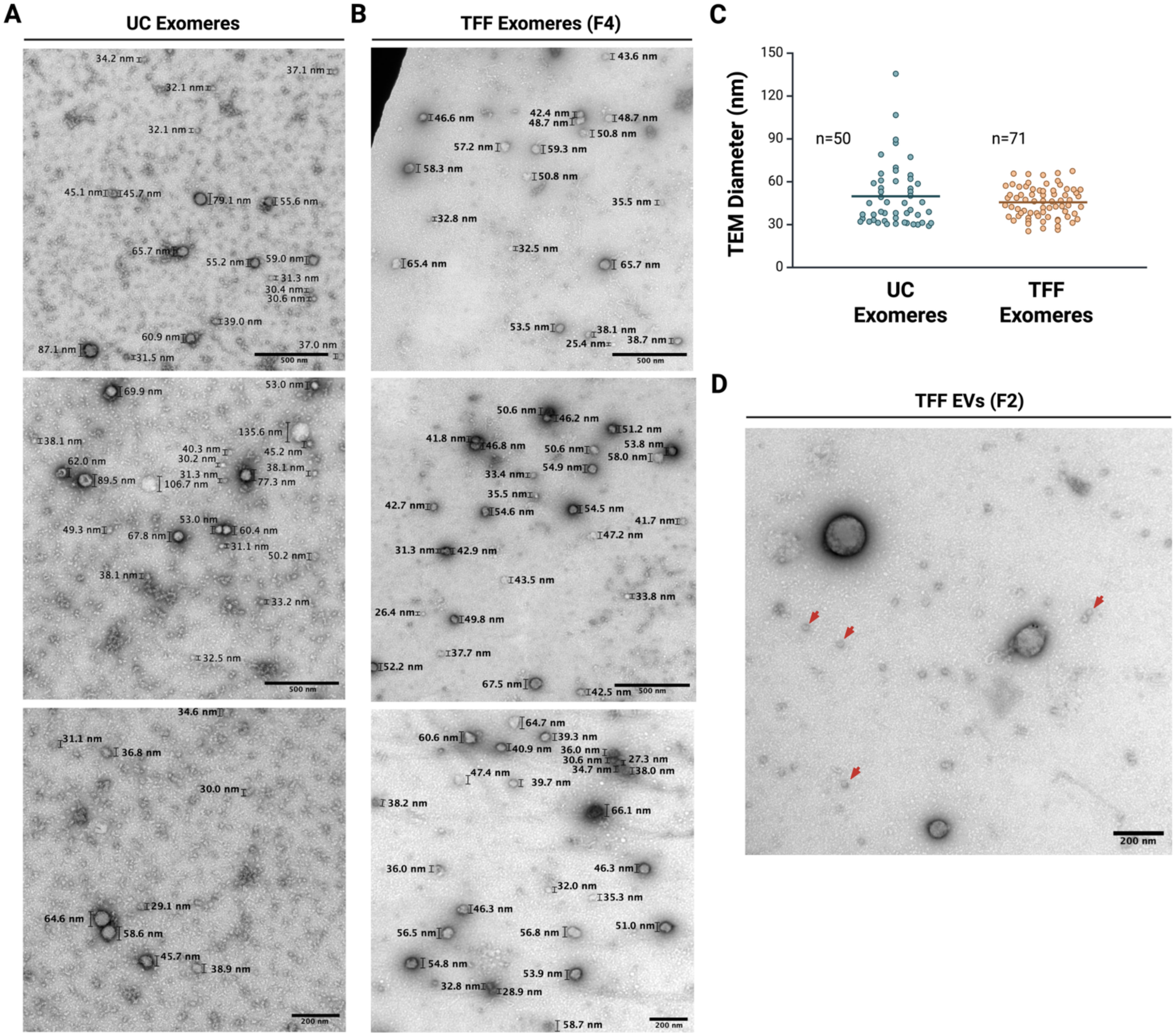


**Supplementary Figure 3:** TEM micrographs of particles measured for **A.** UC exomeres and **B.** TFF exomeres (F4). **C.** Size distribution of particles within each fraction based on manual measurement using ImageJ 1.54g (Java 1.8.0_345). **D.** Exomere-shaped particles co-purify in TFF EV fraction F2. Figure created with BioRender.com.

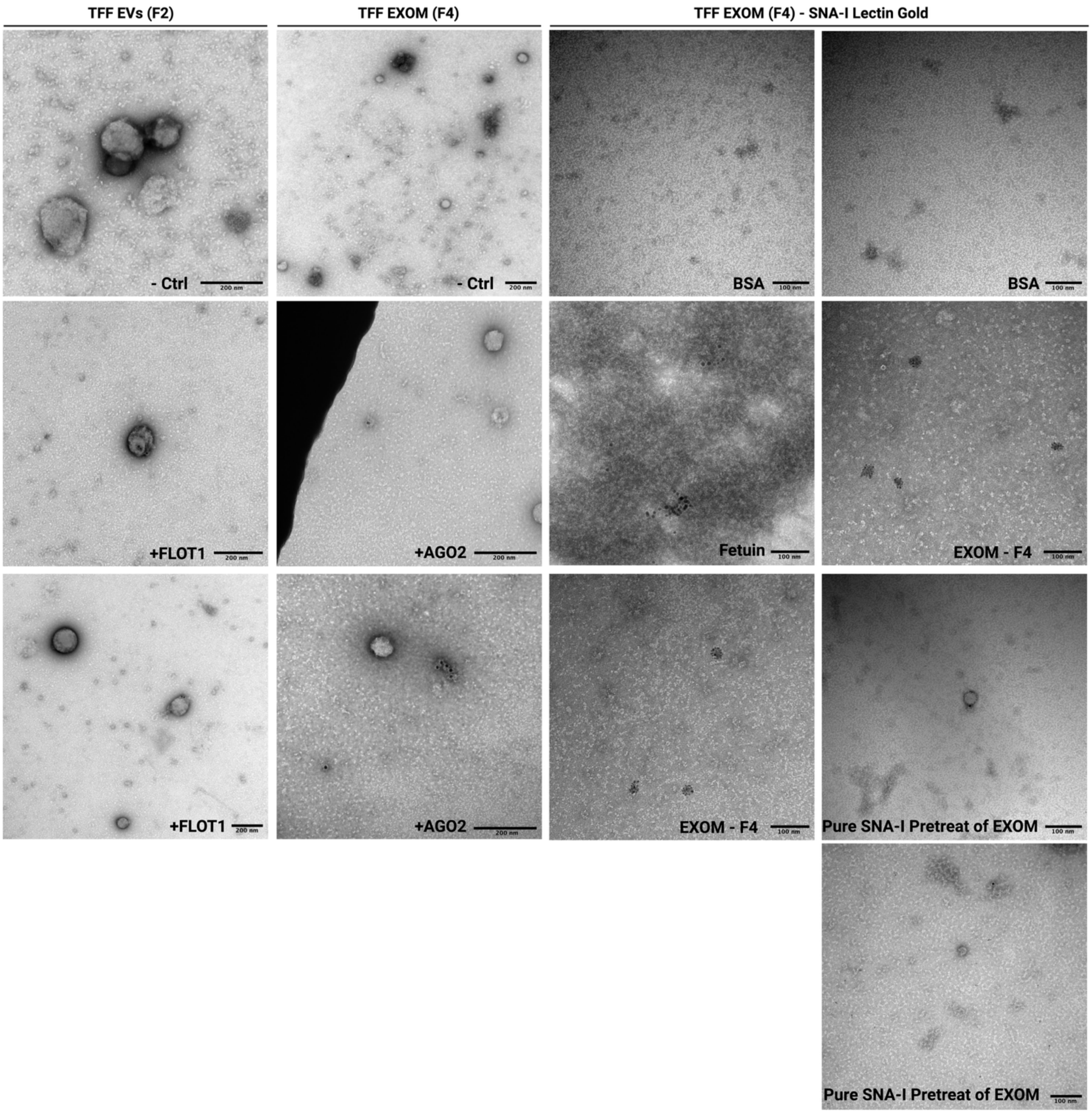


**Supplementary Figure 4:** Unprocessed TEM micrographs for immunogold and SNA-I lectin gold experiments.

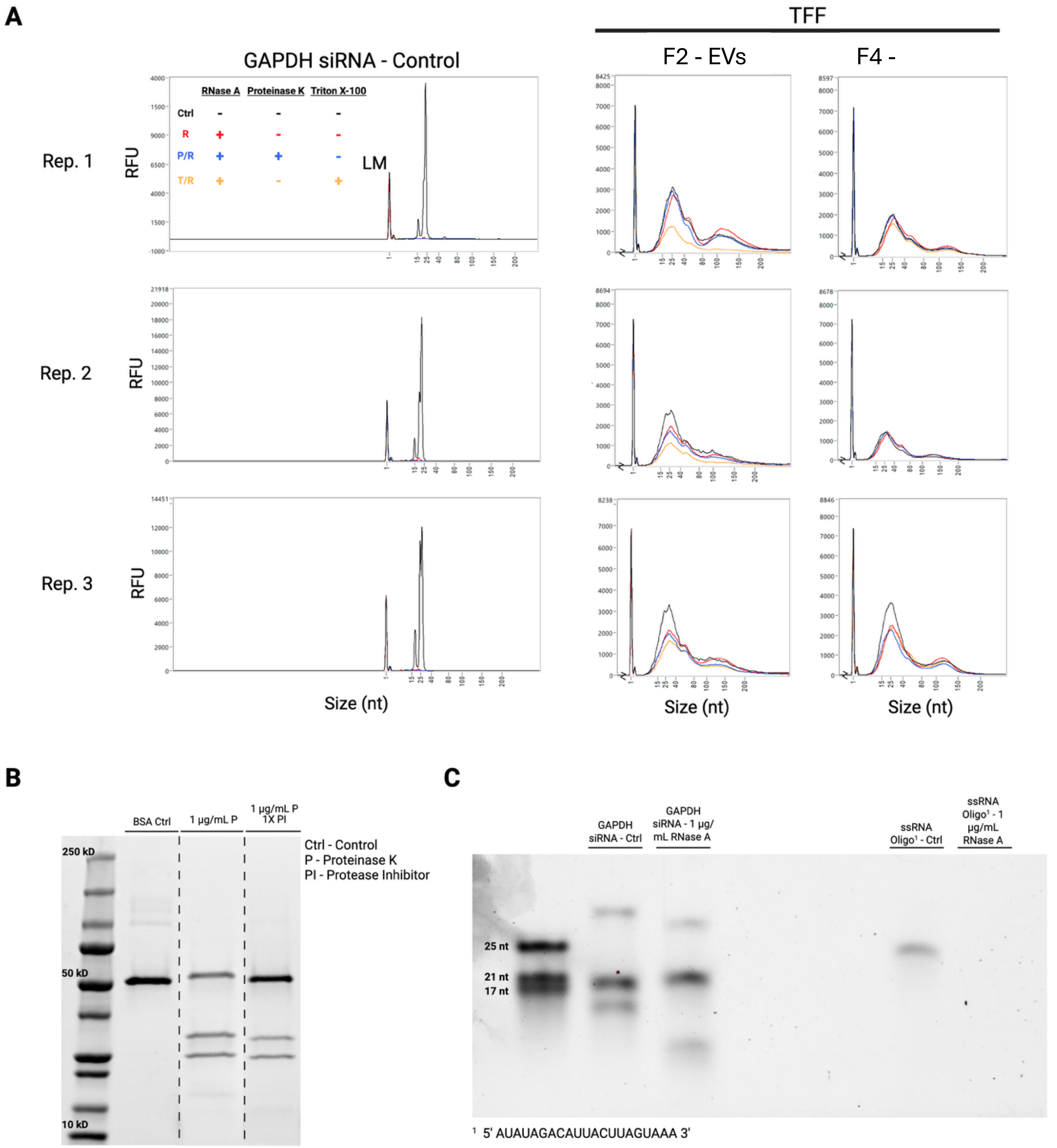


**Supplementary Figure 5: A.** RNA protection assay results representing three biological replicates. Treatment conditions: Ctrl (Control), R (RNase A only), P/R (Proteinase K/RNase A), T/R (Triton X-100/RNase A). **B**. Coomassie blue-stained PAGE gel of BSA treated with Proteinase K or Proteinase K and inhibitor. Gel was cropped to remove conditions not used in the RNase protection experiment. **C.** SYBR$^{TM}$ gold-stained PAGE gel of dsRNA and ssRNA treated with RNase A.

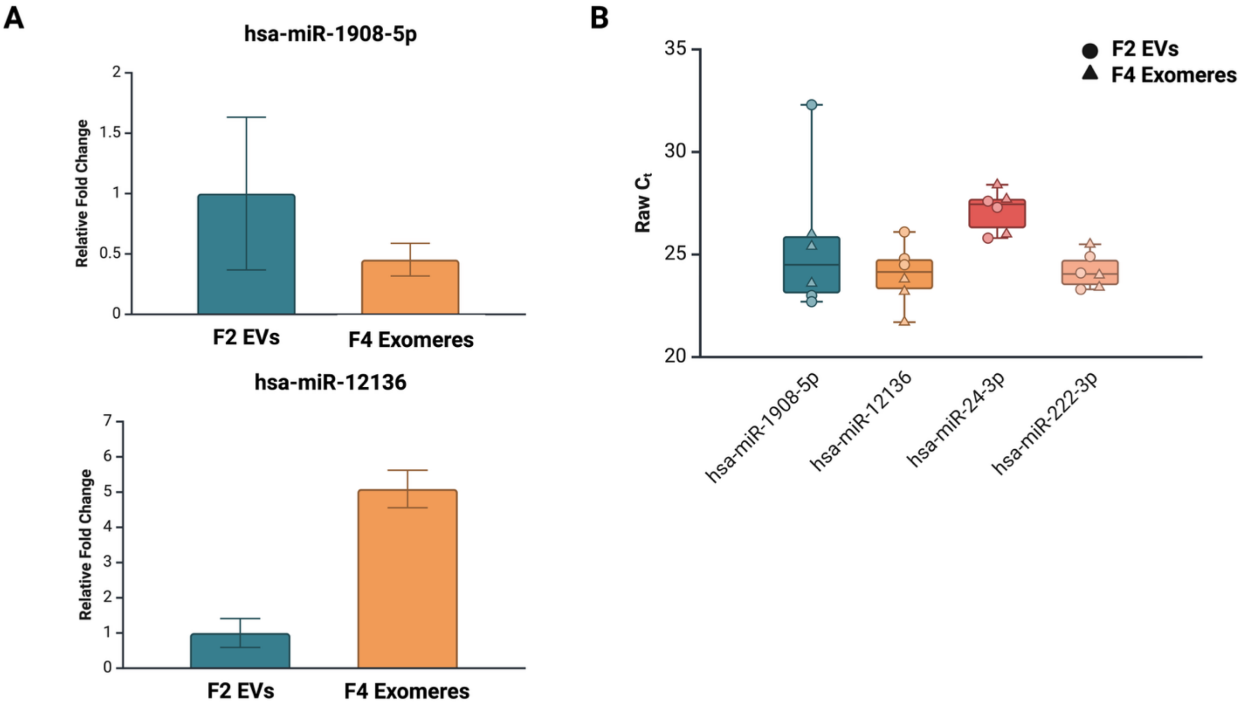


C

| Normalized Read Counts (Median of Ratios) | | | | | | | |
|---|---|---|---|---|---|---|---|
| **miR** | **F2 EVs - 1** | **F2 EVs - 2** | **F2 EVs - 3** | **F4 EXOM - 1** | **F4 EXOM - 2** | **F4 EXOM - 3** | **baseMean** |
| hsa-miR-1908-5p | 7188.83 | 3132.79 | 11269.26 | 298.90 | 541.99 | 375.06 | 3801.14 |
| hsa-miR-12136 | 74.93 | 122.28 | 125.30 | 214.87 | 538.92 | 494.58 | 261.81 |
| hsa-miR-24-3p | 194.82 | 127.17 | 29.48 | 121.24 | 127.38 | 83.20 | 113.88 |
| hsa-miR-222-3p | 1147.94 | 1261.92 | 978.41 | 1162.58 | 1391.40 | 1170.08 | 1185.39 |

**Supplementary Figure 6:** RT-qPCR validation of differentially and stably expressed miRNAs. **A.** Relative fold change calculated using the $\Delta\Delta C_t$ method. F2 EVs were taken as the basis. The geometric mean of hsa-miR-24-3p and hsa-miR-222-3p was used for normalization of each sample. **B.** Raw $C_t$ values for DE miRs and putative reference miRs. **C.** Normalized mean read counts from RNA sequencing for the selected miRs. Figure created with BioRender.com.

| Sample | Reference miRs $C_t$ | | | Sample | DE miRs from RNA Seq $C_t$ | |
|---|---|---|---|---|---|---|
| | hsa-miR-24-3p | hsa-miR-222-3p | Geo Mean $C_t$ | | hsa-miR-1908-5p | hsa-miR-12136 |
| F2 EVs - 1 | 27.3 | 24.1 | 25.7 | F2 EVs - 1 | 32.3 | 26.1 |
| F2 EVs - 2 | 27.6 | 24.9 | 26.2 | F2 EVs - 2 | 23.0 | 24.8 |
| F2 EVs - 3 | 25.8 | 23.3 | 24.5 | F2 EVs - 3 | 22.7 | 24.5 |
| F4 Exomeres - 1 | 28.4 | 25.5 | 26.9 | F4 Exomeres - 1 | 25.4 | 23.8 |
| F4 Exomeres - 2 | 27.7 | 24.0 | 25.8 | F4 Exomeres - 2 | 26.0 | 23.2 |
| F4 Exomeres - 3 | 26.0 | 23.4 | 24.7 | F4 Exomeres - 3 | 23.6 | 21.7 |

| Sample | $\Delta C_t$ | | Sample | $\Delta\Delta C_t$ (F2 EVs as basis) | |
|---|---|---|---|---|---|
| | hsa-miR-1908-5p | hsa-miR-12136 | | hsa-miR-1908-5p | hsa-miR-12136 |
| F2 EVs - 1 | 6.6 | 0.4 | F2 EVs - 1 | 6.1 | 0.8 |
| F2 EVs - 2 | -3.2 | -1.4 | F2 EVs - 2 | -3.8 | -1.1 |
| F2 EVs - 3 | -1.8 | 0.0 | F2 EVs - 3 | -2.4 | 0.3 |
| F4 Exomeres - 1 | -1.5 | -3.1 | F4 Exomeres - 1 | -2.0 | -2.8 |
| F4 Exomeres - 2 | 0.2 | -2.6 | F4 Exomeres - 2 | -0.3 | -2.3 |
| F4 Exomeres - 3 | -1.1 | -3.0 | F4 Exomeres - 3 | -1.6 | -2.6 |

| | | |
|---|---|---|
| Avg. $\Delta C_t$ (F2 as basis) | 0.5 | -0.3 |

| Sample | $2^{-\Delta\Delta Ct}$ (F2 EVs as basis) | | Sample | Relative Fold Change | |
|---|---|---|---|---|---|
| | hsa-miR-1908-5p | hsa-miR-12136 | | hsa-miR-1908-5p | hsa-miR-12136 |
| F2 EVs - 1 | 0.01447 | 0.58329 | F2 EVs - 1 | 0.00233 | 0.49783 |
| F2 EVs - 2 | 13.49239 | 2.12492 | F2 EVs - 2 | 2.17271 | 1.81358 |
| F2 EVs - 3 | 5.12293 | 0.80681 | F2 EVs - 3 | 0.82496 | 0.68860 |
| F4 Exomeres - 1 | 4.14042 | 6.88334 | F4 Exomeres - 1 | 0.66674 | 5.87479 |
| F4 Exomeres - 2 | 1.25052 | 4.77619 | F4 Exomeres - 2 | 0.20137 | 4.07638 |
| F4 Exomeres - 3 | 3.04107 | 6.22430 | F4 Exomeres - 3 | 0.48971 | 5.31231 |

| | | |
|---|---|---|
| Average $2^{-\Delta\Delta Ct}$ (F2 EVs as basis) | 6.20993 | 1.17167 |

**Supplementary Figure 6 (continued):** Relative fold change calculations for F2 EV and F4 exomere miRs validated with RT-qPCR.

**References:**

1. J. Ren, Z. Li and F.-S. Wong, *Journal of Membrane Science*, 2006, **279**, 558–569.

2. S. Singh, K. C. Khulbe, T. Matsuura and P. Ramamurthy, *Journal of Membrane Science*, 1998, **142**, 111–127.
